# The mammalian kinetochore independently regulates its passive and active force-generating interfaces with microtubules

**DOI:** 10.1101/105916

**Authors:** Alexandra F. Long, Dylan B. Udy, Sophie Dumont

## Abstract

The kinetochore links chromosomes to dynamic spindle microtubules and drives both chromosome congression and segregation. To do so, the kinetochore must hold on to depolymerizing and polymerizing microtubules. At metaphase, one sister kinetochore couples to depolymerizing microtubules, pulling its sister along polymerizing microtubules [1,2]. Distinct kinetochore-microtubule interfaces mediate these behaviors: active interfaces transduce microtubule depolymerization into mechanical work, and passive interfaces generate friction as the kinetochore slides along microtubules [3,4]. We do not know the physical and molecular nature [5–7] of these interfaces, or how they are regulated to support diverse mitotic functions in mammalian cells. To address this question, we focus on the mechanical role of the essential load-bearing protein Hec1 [8–11]. Hec1’s affinity for microtubules is regulated by Aurora B phosphorylation on its N-terminal tail [12–15], but its role at the passive and active interfaces remains unclear. Here, we use laser ablation to trigger cellular pulling on mutant kinetochores and decouple sisters *in vivo*, and thereby separately probe Hec1’s role as it moves on polymerizing versus depolymerizing microtubules. We show that Hec1 phosphorylation tunes passive friction along polymerizing microtubules, modulating both the magnitude and timescale of responses to force. In contrast, we find that Hec1 phosphorylation does not affect the kinetochore’s ability to grip depolymerizing microtubules, or switch to this active force-generating state. Together, the data suggest that different kinetochore interfaces engage with growing and shrinking microtubules, and that passive friction can be regulated without disrupting active force generation. Through this mechanism, the kinetochore can modulate its grip on microtubules as its functional needs change during mitosis, and yet retain its ability to couple to microtubules powering chromosome movement.

## Results

### Targeted control of cellular pulling forces on kinetochores in vivo

To probe Hec1’s mechanical role at the mammalian kinetochore-microtubule interface, we sought the ability to exert force on a given kinetochore inside a cell at a specific time. This is necessary to probe the magnitude and timescale of a kinetochore’s response to force, and to perturb kinetochores moving on microtubules in a given polymerization state. We accomplished this using targeted laser ablation to sever one kinetochore-fiber (k-fiber) at metaphase (Fig. 1A).

**Figure 1.**
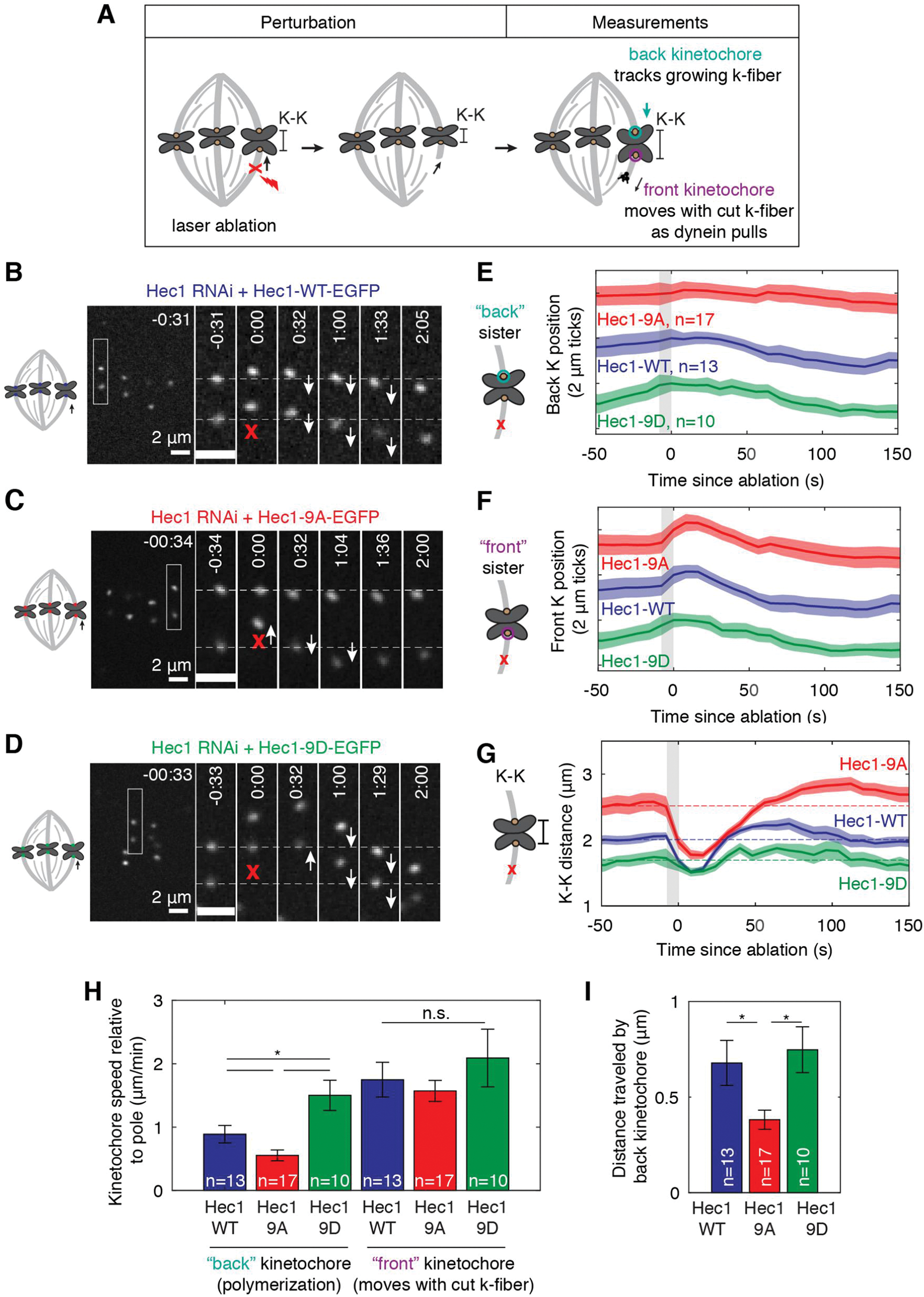
Hec1 tail phosphorylation regulates the magnitude and timescale of the mammalian kinetochore-microtubule interface’s response to force. **(A)** Assay to sever a k-fiber using laser ablation (red X) to induce a dynein-based poleward pulling force on a specific kinetochore pair to probe the back kinetochore’s response to force on polymerizing microtubules. **(B-D)** Timelapse showing representative response of PtK2 **(B)** Hec1-WT-EGFP, **(C)** Hec1-9A-EGFP and **(D)** Hec1-9D-EGFP (each in Hec1 RNAi background) kinetochore pairs to k-fiber laser ablation. First frame after ablation set to 0:00. **(E-G)** Mean positions of Hec1-WT, Hec1-9A, and Hec1-9D **(E)** back and **(F)** front kinetochores and **(G)** K-K distance before and after laser ablation. Kinetochore position is shown normalized to its position before ablation. Traces are mean±SEM and are offset vertically for clarity in **(E,F)**. **(H)** Velocity of the front and back kinetochores during poleward motion (after direction switch) induced by dynein pulling (* for p<0.05, n.s. not significant, Student’s T-test). **(I)** Distance traveled by the back kinetochore over the first 30s of poleward motion after ablation (* for p<0.05, Student’s T-test).

The newly created k-fiber minus-ends recruit dynein, which in turn exerts a poleward pulling force on the attached kinetochore and its sister [16,17].

As a starting point for our Hec1 studies, we expressed Hec1-EGFP in PtK2 cells depleted of endogenous Hec1 by RNAi [11]. We selectively severed polymerizing k-fibers near their kinetochore, and examined the responses of both the “front” and “back” sister kinetochores (proximal and distal to the cut, respectively) (Fig. 1A,B; Movie S1). The response to laser ablation appeared the same as in wild type cells [16,17], and had two phases (Fig. 1B,E-G blue traces; Table 1; n=13). First, the front kinetochore recoiled immediately after cut, reflecting a decrease in force and causing the interkinetochore (K-K) distance to decrease. Second, dynein pulled the microtubules bound to the front kinetochore, moving the sister pair toward the ablation site and increasing the K-K distance. Dynein pulled the front sister faster than its k-fiber polymerized or depolymerized, and faster than normal metaphase movements [16] (Table 1). The front kinetochore’s velocity during dynein pulling was similar between experiments (Fig. 1F,H blue traces; Table 1), consistent with ablation triggering a consistent response. Dynein pulling caused the back kinetochore to turn around within seconds, ultimately pulling it away from its pole along polymerizing microtubules. The back kinetochore’s velocity reported on its ability to passively slide on growing microtubules under force (Fig. 1E,H blue traces; Table1). Thus, targeted k-fiber ablation can produce a pulling force to probe the mechanics of the interface between kinetochores and polymerizing microtubules.

**Table 1.** Role of Hec1 tail phosphorylation in regulating the mechanics of the mammalian kinetochore-microtubule interface.

| Assay | Measurement | Experimental Condition |  |  | T-test<sup>a</sup> |
| --- | --- | --- | --- | --- | --- |
|  |  | Hec1-WT | Hec1-9A | Hec1-9D |  |
| **K-fiber ablation (Fig. 1)**<br>[back kinetochore reflects passive interface] | Number of kinetochores (cells) | 13 (4) | 17 (8) | 10 (7) |  |
|  | K-K distance before ablation (μm) | 2.0 ± 0.1 | 2.5 ± 0.1 | 1.4 ± 0.1 | WT,9A*<br>WT,9D*<br>9A,9D* |
|  | Time at max K-K distance after ablation (s) | 95 ± 7 | 47 ± 5 | n.a.<sup>b</sup> | WT,9A* |
|  | Front kinetochore speed during poleward movement (μm/min) | 1.7 ± 0.3 | 1.6 ± 0.2 | 1.8 ± 0.2 | n.s. |
|  | Distance back kinetochore moves during first 30 s of dynein pulling (μm) | 0.7 ± 0.1 | 0.4 ± 0.1 | 1.0 ± 0.2 | WT,9A*<br>9A,9D* |
|  | Back kinetochore speed during poleward movement (μm/min) | 0.9 ± 0.1 | 0.6 ± 0.1 | 1.7 ± 0.4 | WT,9A*<br>WT,9D*<br>9D,9A* |
| **Metaphase spindle photomarking (Fig. 2)**<br>[passive interface] | Number of kinetochores (cells) | 57 (29) | 60 (24) | 27 (8) |  |
|  | Poleward microtubule flux (μm/min) | 0.65 ± 0.05 | 0.50 ± 0.03 | 0.73 ± 0.07 | WT,9A*<br>9D,9A* |
|  | Kinetochore velocity with respect to microtubule lattice (μm/min) | 0.76 ± 0.04<sup>c</sup> | 0.51 ± 0.03<sup>c</sup> | n.a.<sup>c</sup> | WT,9A* |
|  | Kinetochore velocity > 0 (towards plus-end) with respect to microtubule lattice (μm/min) | 1.25 ± 0.03<sup>c</sup> | 0.83 ± 0.03<sup>c</sup> | n.a.<sup>c</sup> | WT,9A* |
| **Kinetochore ablation (Fig. 3)**<br>[active + passive interface] | Number of kinetochores (cells) | 17 (14) | 21 (12) | 18 (13) |  |
|  | K-K distance before ablation (μm) | 2.3 ± 0.1 | 2.6 ± 0.1 | 1.7 ± 0.1 | WT,9A*<br>WT,9D*<br>9A,9D* |
|  | Poleward kinetochore velocity after sister ablation (μm/min) | 2.0 ± 0.2 | 2.1 ± 0.2 | 2.3 ± 0.2 | n.s. |
|  | Poleward kinetochore velocity relative to microtubule lattice after sister ablation (μm/min)<sup>d</sup> | 1.5 ± 0.2 | 1.4 ± 0.2 | 1.6 ± 0.2 | n.s. |
|  | Time between front sister ablation and back kinetochore switch (s) | 5 ± 3 | 2 ± 2 | 4 ± 2 | n.s. |

Data are presented as mean ± SEM. See also Figures 1-3. n.a. for not applicable. n.s. for not significant.
<sup>a</sup> $p < 0.05$ used as threshold for statistical significance using two tailed Student's T-Test. The abbreviations in the T-test column indicate which of the condition pairs are significantly different (e.g. WT,9D \* indicates a significant difference between Hec1-WT and Hec1-9D).
<sup>b</sup> There is no meaningful maximum K-K distance after ablation for Hec1-9D due to the variability of traces and lack of overshoot above baseline K-K distance from before ablation.
<sup>c</sup> We made these calculations only on the subset of the data collected using photoactivation (Hec1-WT, n=42 and Hec1-9A n=34 kinetochores) since it allowed longer tracking of oscillations. We did not measure kinetochore velocities in Hec1-9D spindles since we were not able to track photomarks for long enough of the kinetochore oscillation cycle.
<sup>d</sup> To adjust velocities to be relative to the microtubule lattice, we assumed poleward flux was unchanged from metaphase measurements (Fig. 2).

### Hec1 tail phosphorylation regulates the magnitude and timescale of the mammalian kinetochore-microtubule interface’s response to force

To probe the mechanical regulation conferred by Hec1’s N-terminal tail phosphorylation during mitosis, we asked whether and how it controls the response of a kinetochore to force. We depleted endogenous Hec1 by RNAi, and expressed either Hec1-9A-EGFP or Hec1-9D-EGFP to mimic constitutive dephosphorylation and phosphorylation, respectively, a range that includes typical Hec1 phosphorylation by Aurora B during mitosis [14]. As expected [14], Hec1-9D and Hec1-9A kinetochores resulted in different steady-state K-K distances (Fig. 1C,D,G; Table 1).

We subjected these Hec1-9A (Fig. 1E-G red traces; n=17; Movie S2) and Hec1-9D kinetochores (Fig. 1E-G, green traces; n=10; Movie S3) to the same force signature as Hec1-WT, as suggested by similar front kinetochore velocities during dynein pulling (Fig. 1H,Table 1). As with Hec1-WT, after k-fiber ablation the front kinetochore recoiled and the K-K distance decreased in both Hec1-9A and Hec1-9D cells. When dynein pulling engaged, however, the back sister responses were different from Hec1-WT. In Hec1-9A cells, the back kinetochore moved more slowly than its front sister (0.6±0.1 vs 1.6±0.2µm/min; Table 1), and moved less far on polymerizing microtubules than Hec1-WT (0.4±0.1 vs 0.7±0.1µm; Fig. 1H,I; Table 1). These differences led to a larger, and longer-lasting, increase in K-K distance above baseline during dynein pulling compared to Hec1-WT (maximum K-K distance was at 95±7 vs 47±5s; Fig. 1G; Table 1). While Hec1-9A kinetochores have more k-fiber microtubules and this could lead to higher force [14,18] on these kinetochores and a larger response, Hec1-9A kinetochores in fact move less. In contrast, in Hec1-9D cells the back sister followed at a rate similar to its front sister (1.7±0.4 vs 1.8±0.2µm/min), which is faster than Hec1-WT (0.9±0.1µm/min), and moved farther than Hec1-WT on polymerizing microtubules (1.0±0.2 vs 0.7±0.1 µm; Fig. 1H,I; Table 1). These responses led to little overshoot in K-K distance above baseline during dynein pulling (Fig. 1 G).

Dephosphorylating Hec1 makes the back kinetochore less mobile in response to force: the back kinetochore moves more slowly and a shorter distance, and takes longer to recover, despite being under higher forces. Phosphorylating Hec1 has the opposite consequences. Thus, Hec1 phosphorylation controls both the magnitude and timescale of the back kinetochore’s response to spindle forces, and thereby sets the effective elasticity and viscosity of the spindle’s reorganization in response to force. Since the back kinetochore’s ability to slide on polymerizing microtubules under force varied depending on the Hec1 tail phosphorylation state, we hypothesized that Hec1’s tail forms part of a passive frictional interface at kinetochores bound to polymerizing microtubules.

### Hec1 tail phosphorylation regulates friction at the passive interface between mammalian kinetochores and polymerizing microtubules

Perturbing Hec1 phosphoregulation is known to change how metaphase sister kinetochores move [13,14], but because sisters are attached it is not clear whether this occurs because of changes in the passive or active interface between the kinetochore and microtubules. To test the hypothesis that Hec1’s tail is part of the passive interface of kinetochores with polymerizing microtubules, we measured how Hec1 phosphorylation changes the velocity – and friction coefficient assuming similar forces – between kinetochore and polymerizing microtubules. To determine kinetochore velocity relative to the microtubule lattice, we tracked kinetochores with Hec1-EGFP phosphomutants, and concurrently measured k-fiber poleward flux [19] by either photomarking PA-GFP-tubulin or photobleaching GFP-tubulin (Fig. 2A-C). K-fiber flux velocities were lower in Hec1-9A (0.50±0.03µm/min, n=60) than in Hec1-9D (0.73±0.07µm/min, n=27) or WT cells (0.65±0.05 µm/min, n=57) (Fig. 2D, Table 1). Consistent with Hec1 phosphorylation decreasing friction at the passive interface, kinetochore velocity with respect to the microtubule lattice during polymerization was higher in Hec1-WT (1.20±0.03µm/min, n=720; Movie S4) than in Hec1-9A cells (0.80±0.03µm/min, n=940; Movie S5) (Fig. 2E, Table 1). Thus, the interface remains dynamic and is never locked within the cell’s Hec1 phosphorylation range; the kinetochore (as a “slip clutch” [4]) can always slip to reduce force on the chromosome – and prevent detachment from microtubules [20].

**Figure 2.**
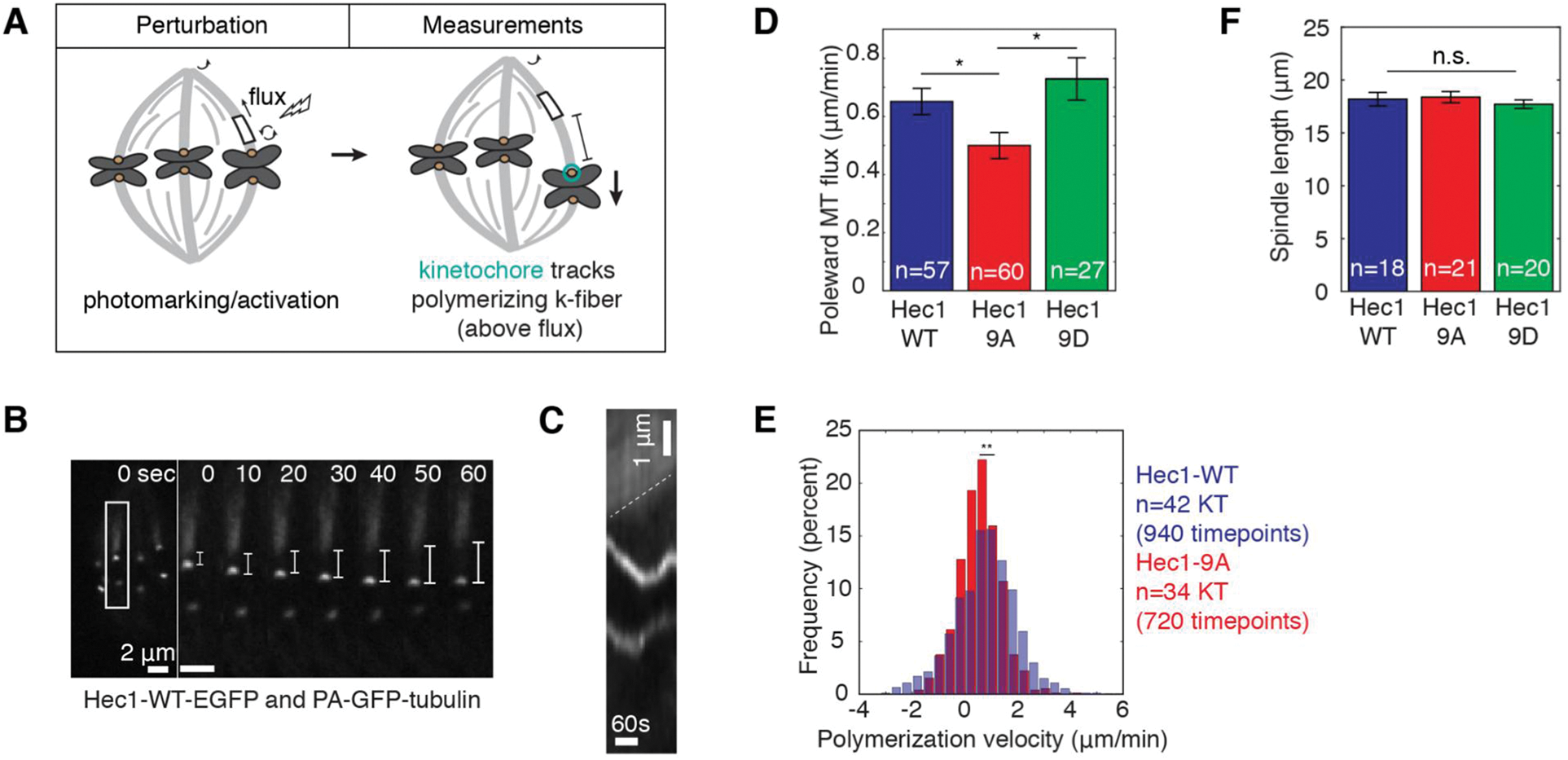
Hec1 tail phosphorylation regulates friction at the passive interface between the mammalian kinetochore and polymerizing microtubules. **(A)** Assay to measure kinetochore velocity relative to the microtubule lattice, tracking kinetochores and measuring poleward k-fiber microtubule flux by photomarking. **(B)** Representative timelapses of Hec1-EGFP and PA-GFP-tubulin PtK2 cells in a Hec1 RNAi background and **(C)** kymograph of poleward microtubule flux (dotted line) measured by photoactivation. Time 0:00 corresponds to photoactivation. The distance between the photomark and the kinetochore (ruler) provides velocity relative to the microtubule lattice. **(D)** Microtubule flux rate (mean±SEM, * for p<0.05, Student’s T-test) in cells with Hec1-WT, Hec1-9A, or Hec1-9D kinetochores (n= number of k-fibers). **(E)** Histogram of kinetochore velocity relative to the microtubule lattice (** for p<0.01, Student’s T-test) **(F)** Spindle length (mean±SEM, n.s. for not significant, p=0.76 one-way ANOVA) in cells with Hec1-WT, Hec1-9A, or Hec1-9D kinetochores (n= number of cells).

These data are consistent with Hec1 being a component of the passive interface of kinetochores with microtubules – whose location was inferred to be in the outer kinetochore [21] Hec1 tail phosphorylation tunes the friction coefficient, and thus the force-velocity relationship, between the mammalian kinetochore and polymerizing microtubules, and does so in a force range relevant to spindle function. Indeed, Hec1 tail phosphorylation has consequences beyond the kinetochore: it changes microtubule flux across the spindle (Fig. 2D), and does so without changing spindle length (Fig. 2F), implying that microtubule depolymerization rates at poles must change. These suggest that Hec1 tail phosphorylation provides a mechanism for regulating spindle mechanics during mitosis, in concert with kinetochore attachment formation.

### Hec1 tail phosphorylation does not disrupt the mammalian kinetochore’s ability to couple to depolymerizing microtubules at the active interface

As we found that Hec1 phosphorylation decreases friction at the passive interface with polymerizing microtubules (Fig. 1-2), we asked whether it also affects the ability to couple to depolymerizing microtubules, thought to require both an active and passive interface [21]. When sister kinetochores are linked (Fig. 1-2), the coupling to depolymerizing microtubules can never be probed directly as it is always resisted by its sister; anaphase kinetochores could in principle provide a solution, but kinetochore biochemistry changes between metaphase and anaphase [22]. Hence, we turned to laser ablation to physically separate sister kinetochores: after ablating one sister, the remaining sister moves towards its pole as its k-fiber depolymerizes [1,23] (Fig. 3A,B).

**Figure 3.**
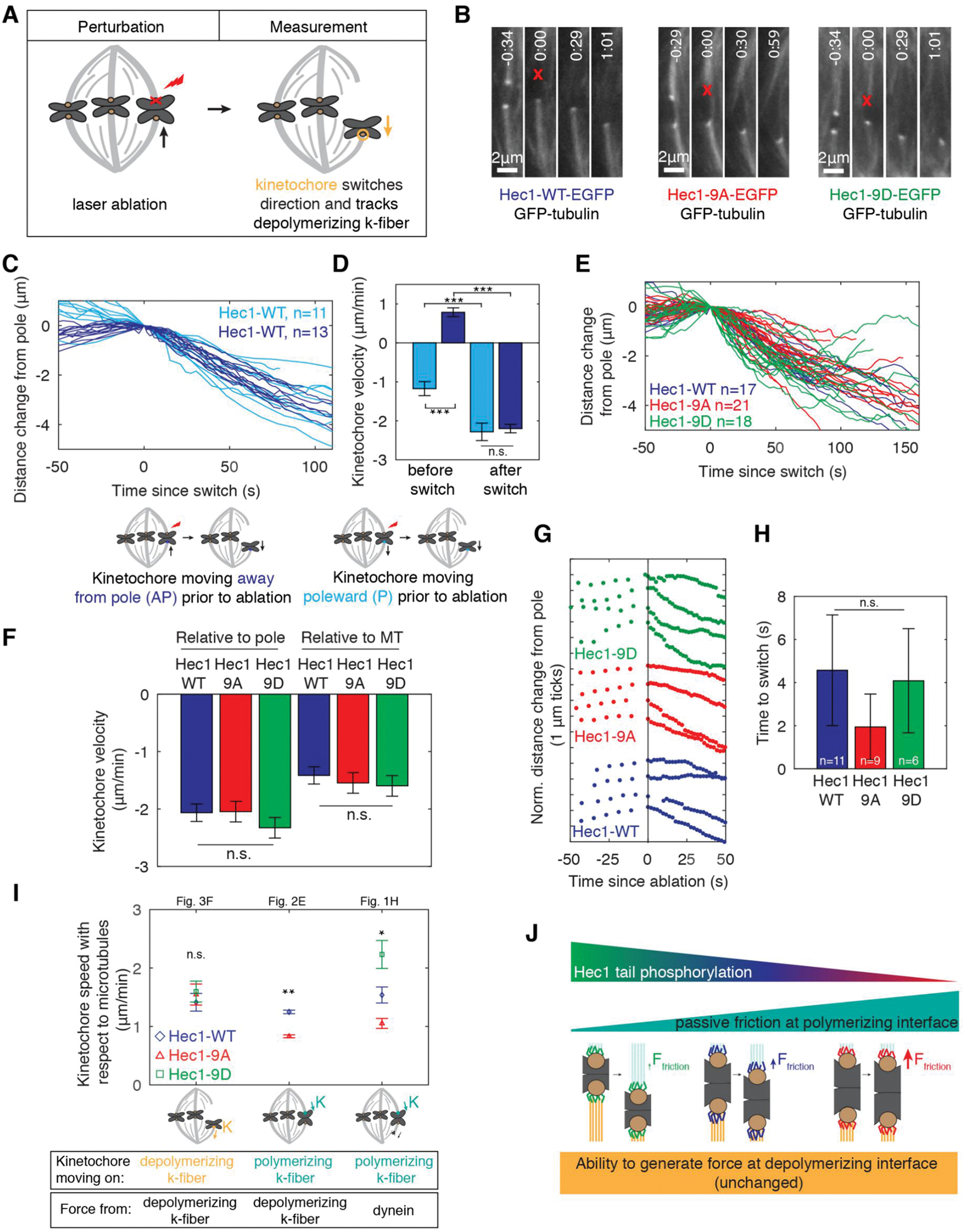
Hec1 tail phosphorylation does not disrupt the mammalian kinetochore’s ability to couple to depolymerizing microtubules at the active interface. **(A)** Assay to decouple sister kinetochores using laser ablation (red X) of one sister kinetochore to probe the remaining sister’s ability to track depolymerizing microtubules. **(B)** Timelapse of Hec1-WT-EGFP, Hec1-9A-EGP, or Hec1-9D-EGFP and GFP-tubulin in PtK2 cells before and after kinetochore ablation. **(C)** Response of kinetochores to sister ablation, with color reflecting the direction prior to ablation (n= number of kinetochores). **(D)** Kinetochore velocity relative to pole before and after its direction switch following sister ablation. (*** for p<0.001, Student’s T-test, n.s. for not significant). **(E)** Responses of kinetochores to sister ablation (n = number of kinetochores). **(F)** Kinetochore velocity after switching to poleward motion (depolymerization) due to ablation of sister. Kinetochore velocities relative to the pole (left) or to the microtubule lattice (right, adjusted for differences in flux from Fig. 2) (same dataset as (D), n.s. for not significant, Student’s T-test). **(G)** Example traces and **(H)** mean delay of kinetochores switching direction after sister ablation (n.s. for not significant, Student’s T-test). **(I)** Summary: Role of Hec1 phosphorylation in regulating kinetochore velocity and function under different mechanical conditions. Kinetochore speeds are replotted from the indicated figures (Fig. 1H values are adjusted for differences in flux from Fig. 2). **(J)** Cartoon summarizing the mechanical role of Hec1 phosphorylation: it regulates friction at the passive interface with polymerizing microtubules (top, cyan) but does not disrupt the mammalian kinetochore’s ability to couple to depolymerizing microtubules at the active interface (bottom, yellow). For simplicity, numbers of microtubules and Hec1 molecules are diagrammed as constant across conditions.

After sister ablation, Hec1-WT kinetochores initially moving poleward speed up, from 1.1±0.2µm/min (depolymerizing microtubules since faster than tubulin flux, Fig. 2D) to 2.3±0.2µm/min (n=11, p<0.01; Fig. 3C,D). This acceleration is consistent with the sister, bound to polymerizing microtubules before ablation, providing resistive friction. In turn, WT kinetochores initially moving away from their pole (polymerizing microtubules) at 0.8±0.1µm/min switch to poleward movement at 2.2±0.2µm/min (n=13; Fig. 3C,D; Movie S6). The directional switch and kinetochore velocity we measure here are faster than those we measured after k-fiber ablation (Fig. 1), which is likely because here there is no resistance from the sister k-fiber interacting with the spindle. Surprisingly, Hec1-9A (Movie S7) and Hec1-9D (Movie S8) kinetochores, which had perturbed K-K distances (Table 1), moved poleward at the same velocity as Hec1-WT after sister ablation (2.0±0.2µm/min, n=21 and 2.3±0.2µm/min, n=18, respectively; Fig. 3B,E,F, Table 1). As kinetochores approach poles, kinetochore velocity remained unchanged despite chromosomes experiencing higher polar ejection forces [23,24]. If there Hec1 phosphorylation regulates the active or passive interface in depolymerization, there is no functional consequence on kinetochore movement in this assay. Hec1-WT, Hec1-9A and Hec1-9D kinetochores also had, within our resolution, indistinguishable times to switch direction from away-from-pole to poleward movement (Fig. 3G,H). This finding suggests that Hec1 phosphorylation does not directly regulate the switch from moving on polymerizing to depolymerizing microtubules. Thus, while Hec1 tail phosphorylation regulates the kinetochore’s ability to passively slide on polymerizing microtubules (Fig. 1-2), it does not affect its ability to couple to and track depolymerizing microtubules or its poleward velocity (Fig. 3).

## Discussion

Accurate chromosome segregation requires the kinetochore to be able to hold on to both polymerizing and depolymerizing microtubules. However, the molecular basis and regulation of the passive and active interfaces that kinetochores use to attach to k-fibers are not known. In particular, separately probing kinetochore movement in defined polymerization states has been challenging. Elegant i*n vitro* assays [25,26] overcome these challenges but are not yet tractable for mammalian kinetochores, while i*n vivo* microneedle [27,28] and laser ablation [1,23,29] studies have probed kinetochore mechanics in defined states, but not their molecular basis. Here, we use a combination of molecular and mechanical perturbations to determine the contribution of Hec1 phosphoregulation to the passive and active interfaces of the mammalian kinetochore. We find that through Hec1 tail phosphorylation, the kinetochore can independently regulate its ability to slide on polymerizing microtubules (passive friction) without losing its ability to couple to depolymerizing microtubules to move chromosomes (active force generation) (Fig. 3I,J). As the needs of mitosis change, regulation of passive friction may set how far and how fast chromosomes move in response to force, and tune whole spindle mechanics, for example increasing mechanical coupling across spindle halves as mitosis progresses.

The basis for Hec1’s tail regulating kinetochore movement along polymerizing but not depolymerizing microtubules is not known. If kinetochore speeds were higher during polymerization than depolymerization states, changes in friction may only be detectable during polymerization; however, we observe higher speeds during depolymerization (Fig. 3I, Table 1). Similarly, direction-specific regulation could in principle arise from differences in microtubule plus-end tip structure, but this structure so far appears not to differ between sisters [30]. Alternatively, Hec1 structure may vary when bound to polymerizing versus depolymerizing microtubules [31,32], or proteins other than Hec1 may bear load and govern chromosome velocity during depolymerization [33–35]. To uncover the molecular basis for Hec1 tail phosphorylation’s direction-dependent role, it will be essential to determine whether such phosphorylation regulates friction directly (by changing the tail’s microtubule affinity) or indirectly (by changing how its other domains, or other proteins, interact with microtubules), and how it does so. Further, it will be essential to map which proteins form the active interface with depolymerizing microtubules.

Our work indicates that Hec1 phosphorylation regulates the mechanics of the mammalian kinetochore-microtubule interface in a direction-dependent manner, revealing a new level of regulation. Hec1 phosphorylation may impact mechanics in both directions *in vitro* when it is the only coupler [36], but only impact mechanics in polymerization *in vivo* due to the presence of – and load-sharing by – other microtubule binding proteins *in vivo*. Consistent with this idea, the Ndc80 tail is nonessential for movement in either direction in budding yeast [37,38], likely because both Ndc80 and the Dam1 complexes bind microtubules [39,40] and provide friction during polymerization, and Dam1 is the main coupler during depolymerization [37]. Functional homologues to Dam1 are being proposed in other eukaryotes [41,42], and the assay we develop here should be helpful in dissecting the mechanical role of these and other proteins in the active and passive microtubule binding interfaces of the mammalian kinetochore. Probing the relative importance of different kinetochore couplers at both interfaces will be critical to understanding the mechanical diversity of kinetochore proteins and functions across systems.

## Author Contributions

Conceptualization, A.F.L, D.B.U., S.D; Methodology, A.F.L., D.B.U., S.D.; Investigation, A.F.L. and D.B.U.; Data Curation, A.F.L; Writing – Original Draft, A.F.L and S.D.; Writing – Review & Editing, A.F.L, D.B.U., S.D.; Visualization, A.F.L.; Funding Acquisition S.D.

## Acknowledgements

We thank Jennifer DeLuca for Hec1-EGFP mutant constructs and advice, Ekaterina Grishchuk, Ronald Vale, David Morgan, and David Agard for discussions, and members of the Dumont Lab for discussions and critical reading of the manuscript. This work was funded by NIH DP2GM119177, NSF CAREER 1554139, the Rita Allen Foundation and Searle Scholars’ Program (S.D.), and a NSF Graduate Research Fellowship (A.F.L.).

## References

1. Khodjakov, A., and Rieder, C.L. (1996). Kinetochores moving away from their associated pole do not exert a significant pushing force on the chromosome. J. Cell Biol. 135, 315–327.

2. Wan, X., Cimini, D., Cameron, L.A., and Salmon, E.D. (2012). The coupling between sister kinetochore directional instability and oscillations in centromere stretch in metaphase PtK1 cells. Mol. Biol. Cell 23, 1035–46.

3. Hill, T.L. (1985). Theoretical problems related to the attachment of microtubules to kinetochores. Proc. Natl. Acad. Sci. U. S. A. 82, 4404–4408.

4. Maddox, P., Straight, A., Coughlin, P., Mitchison, T.J., and Salmon, E.D. (2003). Direct observation of microtubule dynamics at kinetochores in Xenopus extract spindles: implications for spindle mechanics. J. Cell Biol. 162, 377–382.

5. Cheeseman, I.M., and Desai, A. (2008). Molecular architecture of the kinetochore-microtubule interface. Nat. Rev. Mol. Cell Biol. 9, 33–46.

6. Rago, F., and Cheeseman, I.M. (2013). Review series: The functions and consequences of force at kinetochores. J. Cell Biol. 200, 557–565.

7. Sarangapani, K.K., and Asbury, C.L. (2014). Catch and release: how do kinetochores hook the right microtubules during mitosis? Trends Genet. 30, 150–159.

8. DeLuca, J.G., Gall, W.E., Ciferri, C., Cimini, D., Musacchio, A., and Salmon, E.D. (2006). Kinetochore microtubule dynamics and attachment stability are regulated by Hec1. Cell 127, 969–982.

9. Cheeseman, I.M., Chappie, J.S., Wilson-Kubalek, E.M., and Desai, A. (2006). The Conserved KMN Network Constitutes the Core Microtubule-Binding Site of the Kinetochore. Cell 127, 983–997.

10. Ciferri, C., Pasqualato, S., Screpanti, E., Varetti, G., Santaguida, S., Dos Reis, G., Maiolica, A., Polka, J., De Luca, J.G., De Wulf, P., et al. (2008). Implications for kinetochore-microtubule attachment from the structure of an engineered Ndc80 complex. Cell 133, 427–39.

11. Guimaraes, G.J., Dong, Y., McEwen, B.F., and Deluca, J.G. (2008). Kinetochore-microtubule attachment relies on the disordered N-terminal tail domain of Hec1. Curr. Biol. 18, 1778–1784.

12. Powers, A.F., Franck, A.D., Gestaut, D.R., Cooper, J., Gracyzk, B., Wei, R.R., Wordeman, L., Davis, T.N., and Asbury, C.L. (2009). The Ndc80 kinetochore complex forms load-bearing attachments to dynamic microtubule tips via biased diffusion. Cell 136, 865–875.

13. DeLuca, K.F., Lens, S.M.A., and DeLuca, J.G. (2011). Temporal changes in Hec1 phosphorylation control kinetochore-microtubule attachment stability during mitosis. J. Cell Sci. 124, 622–634.

14. Zaytsev, A. V., Sundin, L.J.R., DeLuca, K.F., Grishchuk, E.L., and DeLuca, J.G. (2014). Accurate phosphoregulation of kinetochore-microtubule affinity requires unconstrained molecular interactions. J. Cell Biol. 206, 45–59.

15. Zaytsev, A. V., Mick, J.E., Maslennikov, E., Nikashin, B., DeLuca, J.G., and Grishchuk, E.L. (2015). Multisite phosphorylation of the NDC80 complex gradually tunes its microtubule-binding affinity. Mol. Biol. Cell 26, 1829–1844.

16. Elting, M.W., Hueschen, C.L., Udy, D.B., and Dumont, S. (2014). Force on spindle microtubule minus ends moves chromosomes. J. Cell Biol. 206, 245–256.

17. Sikirzhytski, V., Magidson, V., Steinman, J.B., He, J., Le Berre, M., Tikhonenko, I., Ault, J.G., McEwen, B.F., Chen, J.K., Sui, H., et al. (2014). Direct kinetochore-spindle pole connections are not required for chromosome segregation. J. Cell Biol. 206, 231–243.

18. Cimini, D., Cameron, L.A., and Salmon, E.D. (2004). Anaphase Spindle Mechanics Prevent Mis-Segregation of Merotelically Oriented Chromosomes. Curr. Biol. 14, 2149–2155.

19. Mitchison, T.J. (1989). Polewards Microtubule Flux in the Mitotic Spindle: J. Cell Biol. 109, 637–652.

20. Matos, I., Pereira, A.J., Lince-Faria, M., Cameron, L.A., Salmon, E.D., and Maiato, H. (2009). Synchronizing chromosome segregation by flux-dependent force equalization at kinetochores. J. Cell Biol. 186, 11–26.

21. Dumont, S., Salmon, E.D., and Mitchison, T.J. (2012). Deformations within moving kinetochores reveal different sites of active and passive force generation. Science 337, 355–358.

22. Su, K.-C., Barry, Z., Schweizer, N., Maiato, H., Bathe, M., and Cheeseman, I.M. (2016). A Regulatory Switch Alters Chromosome Motions at the Metaphase-to-Anaphase Transition. Cell Rep. 17, 1728–1738.

23. Skibbens, R. V., Rieder, C.L., and Salmon, E.D. (1995). Kinetochore motility after severing between sister centromeres using laser microsurgery: evidence that kinetochore directional instability and position is regulated by tension. J. Cell Sci. 108, 2537–2548.

24. Ke, K., Cheng, J., and Hunt, A.J. (2009). The Distribution of Polar Ejection Forces Determines the Amplitude of Chromosome Directional Instability. Curr. Biol. 19, 807–815.

25. Akiyoshi, B., Sarangapani, K.K., Powers, A.F., Nelson, C.R., Reichow, S.L., Arellano-Santoyo, H., Gonen, T., Ranish, J.A., Asbury, C.L., and Biggins, S. (2010). Tension directly stabilizes reconstituted kinetochore-microtubule attachments. Nature 468, 576–579.

26. Miller, M.P., Asbury, C.L., and Biggins, S. (2016). A TOG Protein Confers Tension Sensitivity to Kinetochore-Microtubule Attachments. Cell 165, 1428–1439.

27. Nicklas, R.B., and Staehly, C.A. (1967). Chromosome micromanipulation. I. The mechanics of chromosome attachment to the spindle. Chromosoma 21, 1–16.

28. Skibbens, R. V., and Salmon, E.D. (1997). Micromanipulation of chromosomes in mitotic vertebrate tissue cells: tension controls the state of kinetochore movement. Exp. Cell Res. 235, 314–324.

29. McNeill, P.A., and Berns, M.W. (1981). Chromosome behavior after laser microirradiation of a single kinetochore in mitotic PtK2 cells. J. Cell Biol. 88, 543–553.

30. McIntosh, J.R., Grishchuk, E.L., Morphew, M.K., Efremov, A.K., Zhudenkov, K., Volkov, V.A., Cheeseman, I.M., Desai, A., Mastronarde, D.N., and Ataullakhanov, F.I. (2008). Fibrils connect microtubule tips with kinetochores: a mechanism to couple tubulin dynamics to chromosome motion. Cell 135, 322–333.

31. Wang, H.-W., Long, S., Ciferri, C., Westermann, S., Drubin, D., Barnes, G., and Nogales, E. (2008). Architecture and Flexibility of the Yeast Ndc80 Kinetochore Complex.

32. Kudalkar, E.M., Scarborough, E.A., Umbreit, N.T., Zelter, A., Gestaut, D.R., Riffle, M., Johnson, R.S., MacCoss, M.J., Asbury, C.L., and Davis, T.N. (2015). Regulation of outer kinetochore Ndc80 complex-based microtubule attachments by the central kinetochore Mis12/MIND complex. Proc. Natl. Acad. Sci. 112, 5583–5589.

33. Nicklas, R.B. (1983). Measurements of the force produced by the mitotic spindle in anaphase. J. Cell Biol. 97, 542–548.

34. Inoué, S., and Salmon, E.D. (1995). Force generation by microtubule assembly/disassembly in mitosis and related movements. Mol. Biol. Cell 6, 1619–1640.

35. Joglekar, A.P., Bloom, K.S., and Salmon, E.D. (2010). Mechanisms of force generation by end-on kinetochore-microtubule attachments. Curr. Opin. Cell Biol. 22, 57–67.

36. Umbreit, N.T., Gestaut, D.R., Tien, J.F., Vollmar, B.S., Gonen, T., Asbury, C.L., and Davis, T.N. (2012). The Ndc80 kinetochore complex directly modulates microtubule dynamics. Proc. Natl. Acad. Sci. U. S. A. 109, 16113–16118.

37. Suzuki, A., Badger, B.L., Haase, J., Ohashi, T., Erickson, H.P., Salmon, E.D., and Bloom, K.S. (2016). How the kinetochore harnesses microtubule force and centromere stretch to move chromosomes revealed by a FRET tension sensor within Ndc80 protein. Nat. Cell Biol. 18, 382–392.

38. Sarangapani, K.K., Akiyoshi, B., Duggan, N.M., Biggins, S., and Asbury, C.L. (2013). Phosphoregulation promotes release of kinetochores from dynamic microtubules via multiple mechanisms. Proc. Natl. Acad. Sci. U. S. A. 110, 7282–7287.

39. Lampert, F., Hornung, P., and Westermann, S. (2010). The Dam1 complex confers microtubule plus end-tracking activity to the Ndc80 kinetochore complex. J. Cell Biol. 189, 641–649.

40. Tien, J.F., Umbreit, N.T., Gestaut, D.R., Franck, A.D., Cooper, J., Wordeman, L., Gonen, T., Asbury, C.L., and Davis, T.N. (2010). Cooperation of the Dam1 and Ndc80 kinetochore complexes enhances microtubule coupling and is regulated by aurora B. J. Cell Biol. 189, 713–723.

41. Schmidt, J.C., Arthanari, H., Boeszoermenyi, A., Dashkevich, N.M., Wilson-Kubalek, E.M., Monnier, N., Markus, M., Oberer, M., Milligan, R.A., Bathe, M., et al. (2012). The Kinetochore-Bound Ska1 Complex Tracks Depolymerizing Microtubules and Binds to Curved Protofilaments. Dev. Cell 23, 968–980.

42. Hanisch, A., Silljé, H.H.W., and Nigg, E.A. (2006). Timely anaphase onset requires a novel spindle and kinetochore complex comprising Ska1 and Ska2. EMBO J. 25, 5504–5515.

